# CRISPR Cas9 searches for a protospacer adjacent motif by one-dimensional diffusion

**DOI:** 10.1101/264879

**Authors:** Viktorija Globyte, Seung Hwan Lee, Taegun Bae, Jin-Soo Kim, Chirlmin Joo

## Abstract

Since its discovery, the CRISPR/Cas9 system has been at the focus of fundamental researchers, genome engineers, and the general public alike. Despite being in the spotlight for several years, aspects of the precise molecular mechanism of Cas9 activity remain ambiguous. We use single-molecule Foerster resonance energy transfer (smFRET) to reveal Cas9 target search mechanism with nanometer sensitivity. We have developed single-molecule assays to monitor transient interactions of Cas9 and DNA in real time. Our study shows that Cas9 interacts with the protospacer adjacent motif (PAM) sequence weakly, yet probing neighboring sequences via lateral diffusion. This dynamic mode of interactions leads to translocation of Cas9 to another PAM nearby and consequently an on-target sequence. We propose a model in which lateral diffusion competes with 3-dimensional diffusion and thus might aid PAM finding and consequently on-target binding.

## Introduction

CRISPR (Clustered Regularly Interspaced Short Palindromic Repeats)-Cas systems are adaptive prokaryotic immune systems that provide bacteria and archea with a defense mechanism against invading foreign genetic elements^1–6^. Upon infection, fragments of the invader’s DNA are incorporated into the CRISPR locus in the host genome ^7–10^. Those fragments are then transcribed into short CRISPR RNAs (crRNAs) which assemble with CRISPR-associated (Cas) proteins in order to recognize and destroy the invader when it returns ^11^. The most famous of the discovered CRISPR systems is the type II system where the DNA of the invader is recognized and destroyed by the Cas9 protein which assembles with two RNA molecules, namely crRNA and trans-activating crRNA (tracrRNA) ^4,5,10,12–14^. The most widely researched Cas9 ortholog, *Streptococcus pyogenes* Cas9, recognizes a 20-nt target which is flanked by a PAM (protospacer adjacent motif) sequence on the 3‘ end of the target ^12,13^. The PAM sequence for SpCas9 is 5‘-NGG-3‘. CRISPR-Cas9 system has gained enormous attention due to its use in genome editing owing to its simplicity and programmability ^15–18^. In order to edit a gene, Cas9 first has to find a small 23-nt sequence in a genome containing kilo-bases or mega-bases of DNA, in a crowded cellular environment. This process has been demonstrated to be slow, with a single Cas9 protein requiring 6 hours to locate a single target in a bacterial cell ^19^. This example shows that efficient targeting of specific genes requires an advanced knowledge of Cas9 target search mechanism.

A recent single-molecule study using “DNA curtains” has shown that Cas9 uses only 3-dimensional diffusion to locate its target ^20^. Due to diffraction limit (∼100 nm) of the DNA curtain technique, it remains unknown whether the model of exclusive 3-dimensional target search is valid for the length scale of nucleotides. Other single-molecule studies have shown that RNA-guided proteins such as Argonaute or CRISPR type I Cascade protein complex use facilitated 1-dimensional diffusion during their target search ^21,22^ Using single-molecule FRET (Foerster Resonance Energy Transfer), we investigate the target search mechanism of SpCas9 and demonstrate that it uses 1-D diffusion along the DNA strand during its target search.

## Results

### Single-Molecule Observation of Cas9 PAM Search

To visualize Cas9 target search process on a nanometer scale, we used single-molecule Förster resonance energy transfer (smFRET) technique. Biotinylated Cas9 (**Supplementary Fig. 1a**) was immobilized on a PEG-coated quartz surface for long-term observation (Fig. 1a) ^21^. Biotinylation was shown not to affect Cas9 catalytic activity (**Supplementary Fig. 1b**). The Cas9 protein was pre-incubated with dye-labeled crRNA and tracrRNA for 20 minutes before surface immobilization. Free-floating molecules, such as any unattached RNA or Cas9 molecules, were washed away, and remaining immobilized Cas9:RNA complexes could be directly imaged using total internal reflection microscopy (Fig. 1a). Synthetic DNA and crRNA substrates were labeled with Cy3 and Cy5 dyes, respectively, such that the FRET efficiency between them would report on the position where Cas9 localizes on the DNA. Addition of Cy3-labeled DNA substrates did not affect the immobilization of Cas9:RNA complexes (Fig. 1a). When excited with a green laser, binding events appear as spots on the CCD image (Cy3 signal on the left side and Cy5 signal on the right side) (Fig. 1b,c). In fluorescence time traces binding events are characterized by increase in fluorescence intensity due to either direct excitation of donor or energy transfer from donor to acceptor (Fig. 1b,c).

**Figure 1.**
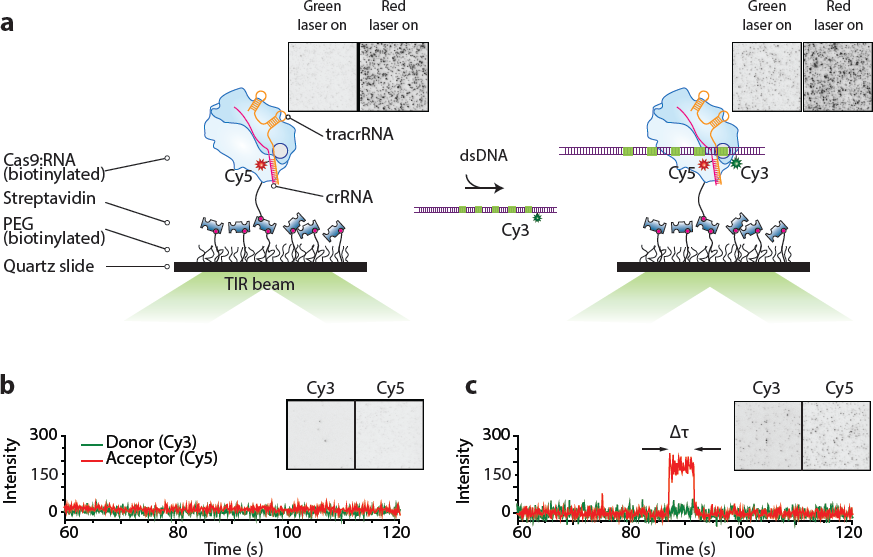
Single-molecule assay for observation of Cas9-DNA interactions. A) Illustrated schematic representation of our single-molecule assay. Cas9 assembled with tracrRNA and Cy5-labelled crRNA is immobilized on the surface via biotin-streptavidin links (left). After addition of Cy3-labelled DNA substrate interaction between Cas9 and DNA brings the dyes together resulting in FRET signal (right). EmCCD images show Cas9:RNA immobilized on the surface with green and red laser illumination (left). No fluorescence is seen when illuminated with green laser. Immobilized Cas9 can be seen when illuminated with red laser. After addition of Cy3-labeled DNA spots appear on the EmCCD in the green channel when illuminated with green laser (right). Size of EmCCD images is 25µm × 25µm.
B) Example time trace when no binding event occurs (green laser excitation). EmCCD image shows signal from green and red channels before the addition of Cy3-labelled DNA when illuminated with green laser. Size of EmCCD image is 25µm × 50 µm.
C) Example time trace showing a binding event (green laser excitation). EmCCD image shows signal from green and red channels after the addition of Cy3-labeled DNA when illuminated with green laser. Size of EmCCD image is 25µm × 50 µm.

The initial step in Cas9 target search is finding and recognizing a PAM sequence. It has been shown that, without encountering a cognate PAM, Cas9 cannot start R-loop formation, despite the full complementarity between the guide RNA and the target ^23^. Therefore, in order to elucidate the mechanism by which Cas9 finds and recognizes PAM alone, we first investigated how the protein interacts with DNA strands containing only PAM sequences when no target sequence is present. In particular, multiple binding sites in close proximity have been shown to cause a synergistic effect in another RNA-guided target search system ^24^. This effect emerges when the interaction between a searcher and a target is characterized by more than simple one-step binding and dissociation ^21,25^. Such a synergistic effect, if observed, would be an indication that Cas9 uses an additional mechanism to 3D diffusion when searching for PAM sequences.

To investigate how Cas9 interacts with multiple PAM sites and whether such synergistic effect exists in the CRISPR-Cas9 system, we designed DNA constructs containing 0, 1, 2, 3, 4 and 5 equidistant PAM sites (Fig. 2a). DNA was labeled with a Cy3 dye at position −8 with respect to the first guanine on the first PAM site on the 5’ end of the target DNA strand (Fig. 2a). crRNA was labeled with a Cy5 dye outside the guide region such that Cas9:RNA complex binding to the first PAM site would yield a high FRET value (Fig. 2 a,b). Binding to other PAM sequences would increase distance between dyes and therefore were expected to yield lower and distinct FRET values for each site, thus allowing to distinguish which PAM site was bound (Fig 2b).

**Figure 2.**
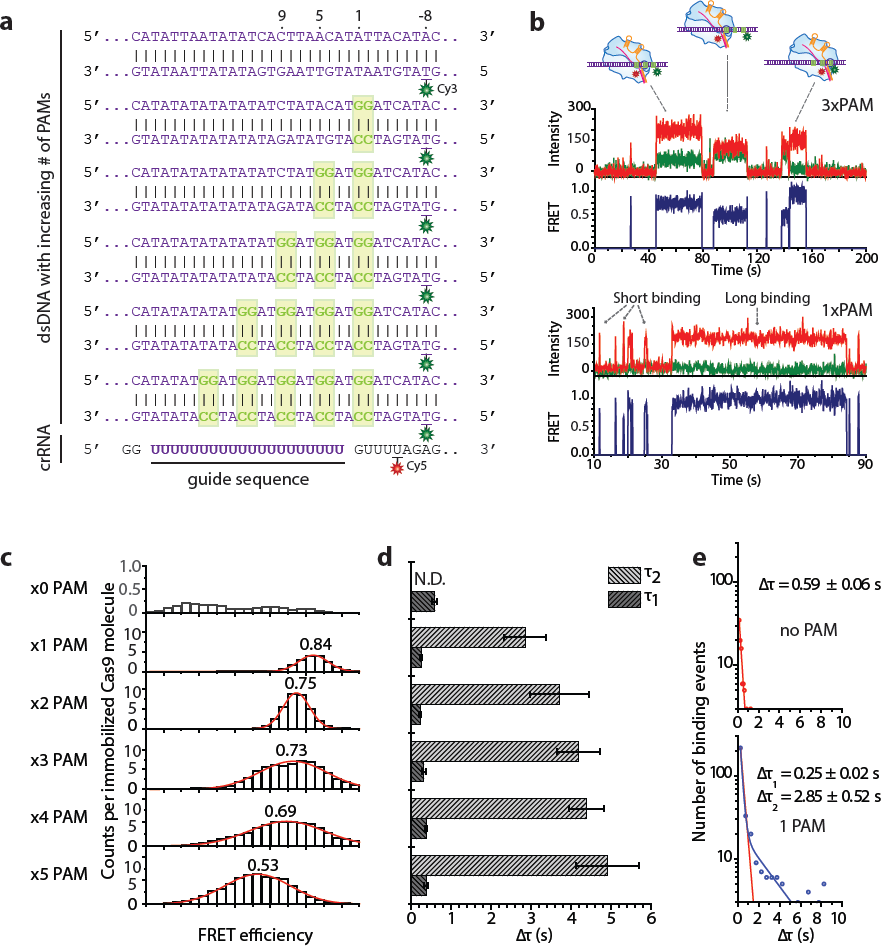
PAM search mechanism. A) DNA and crRNA sequences used for multiple PAM experiments with indicated labeling positions. Di-nucleotide GG PAMs are shown in boxes.
B) Example time traces when DNA containing 3 PAM sequences is added (top) and when DNA containing a single PAM is added showing both short and long binding events (bottom).
C) FRET histograms of all binding events for each construct. Y axis represents total number of counts, divided by the number of immobilized molecules in the field of view.
D) Bar plot of all dwelltime values for all constructs showing short binding events for all DNA and long events only for the constructs containing PAM sequences. The times shown are mean values of dwelltimes obtained during four different experiments on four different days. Error bars represent standard error of the mean.
E) Example dwelltime histograms from negative control and a construct containing a single PAM. Negative control dwelltime distribution is characterized by a single exponential decay (top) and dwelltime distribution for PAM-containing DNA is fitted by a double-exponential decay (blue line) (bottom). Equation for the double exponential decay fit: 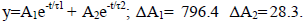 Red line shows what a single-exponential decay fit for such distribution would be. Errors represent standard error of the mean. Dwelltime distributions for remaining DNA constructs are shown in Supplementary Fig. 2c.

Negative control construct containing no PAM sites showed a very low number of binding events with random FRET values (Fig. 2c, **Supplementary Fig. 2a**). Binding to DNA containing a single PAM yielded a narrow FRET distribution centered at 0.84 (Fig. 2c). Binding events showing different FRET states appeared when the number of PAM sites increased (Fig 2b). FRET histogram of all binding events for the construct containing 2 PAM sites shifted to a lower FRET value of 0.75 (Fig. 2c). Lower FRET states appeared for DNA constructs containing 3 and 4 PAMs with FRET histograms broadening significantly and the peaks of the histograms shifting to 0.73 and 0.69 respectively. (Fig. 2c). Finally, a histogram of all binding events for the construct containing 5 PAM sites was broad and centered at 0.53 - an average of all FRET values yielded by specific binding to any of the five PAM sites on the DNA strand (Fig. 2c). Furthermore, multiple binding events showing FRET efficiencies corresponding to different PAMs could be observed in a single FRET trace showing that a single Cas9 can bind and dissociate from different PAM sites, as expected (Fig. 2b).

In addition to broadening of the FRET histograms, the dwelltime was observed to increase with increasing number of PAM sequences (Fig. 2d). Experiments on DNA constructs without any PAM or target sequences only showed short-lived binding (Δτ_1_), with a dwelltime of 0.59±0.06 s (Fig. 2d, e). Short binding events were also observed with PAM-containing DNA constructs ranging between 0.25±0.02 s to 0.37±0.04 s – similar for all constructs, regardless of the number of PAM sequences (Fig. 2d, **Supplementary Fig. 2c**). Interestingly, a second type of longer binding events was observed for the constructs containing PAM sites as opposed to the negative control (Fig. 2b, d, e, **Supplementary Fig. 2c**). It is further noted that the second dweltime (Δτ_2_) increased with increasing number of adjacent PAM sites from 2.85±0.52 s for the construct with a single PAM to 4.91±0.78 s for the construct with 5 PAM sites (Fig. 2d). Further analysis showed no correlation between FRET values and dwelltimes, suggesting that it is not the position of a PAM site, but rather the number of PAM sites in close proximity that caused this increase (**Supplementary Fig. 2a**).

The observation that even for a single PAM the binding time is characterized by a double exponential distribution suggests that Cas9 uses another mechanism in addition to 3-D diffusion during its target search, as processes following exclusively 3-D diffusion follow one-step dissociation kinetics ^26^. Furthermore, the increase of τ_2_ implies that, due to the presence of multiple PAM sites in close proximity, Cas9 experiences a synergistic effect which causes it to stay bound to DNA for longer. We therefore hypothesize that upon encountering a PAM, Cas9 can follow two pathways. First, Cas9 can dissociate from DNA in a 3-dimensional fashion upon failing to form an RNA-DNA R-loop (corresponding to τ_1_). Secondly, Cas9 can locally diffuse in a 1-dimenional fashion probing adjacent PAM sites (corresponding to τ_2_).

### 1-Dimensional Diffusion Used for Target Search

The observation that Cas9 stays bound to a DNA strand for longer when multiple neighboring PAM sequences are present, suggested that this effect could be caused by lateral diffusion between the PAM sites ^26^. To investigate the possibility that Cas9 is able to scan PAM sequences in a 1-D fashion and to explore whether such probing could lead to binding to a neighboring target site, we designed DNA constructs containing a partial target of 9 nt and an increasing number of PAM sites adjacent to it: 1xPAM, 3xPAM, 5xPAM, 7xPAM and 9xPAM (Fig 3a). DNA was labeled at position +13 on the target strand and crRNA at position +10 relative to the first complementary nucleotide, such that high FRET efficiency would only be observed upon productive binding to the partial target site (Fig 3a). Partial complementarity was chosen in order to allow for observation of multiple binding events ^23,27^.

**Figure 3.**
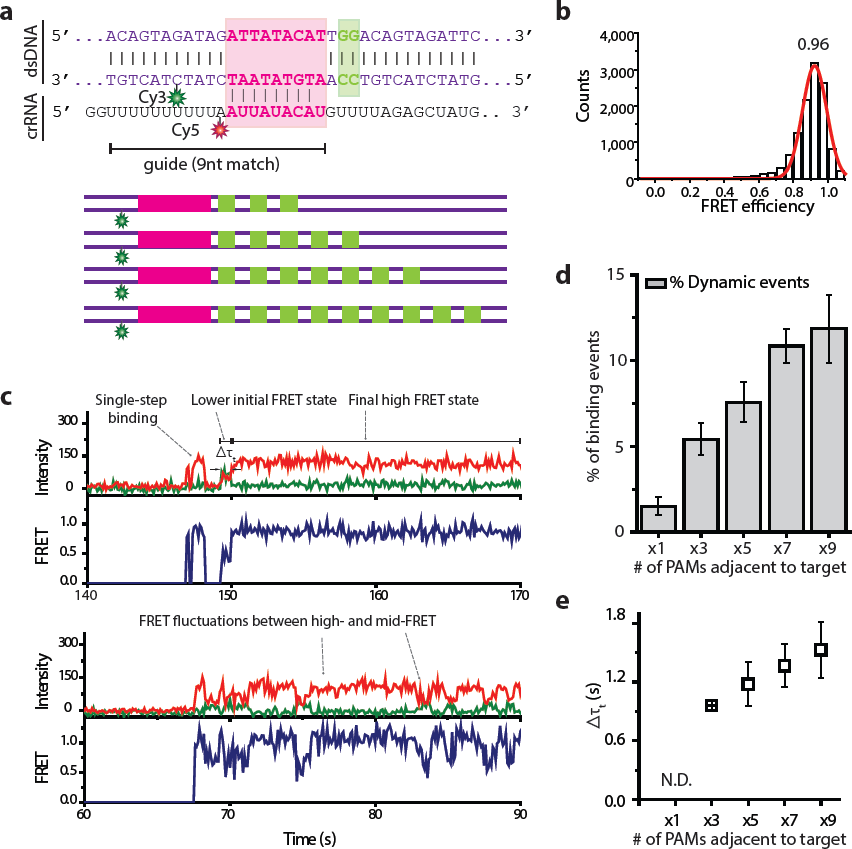
Multiple PAM influence on binding to a target site. A) DNA and crRNA sequence used for experiments of a single target and multiple PAMs. DNA sequence is shown for the single-target single-PAM construct and schematic representation is shown for multiple-PAM containing constructs.
B) FRET histogram of all binding events from the construct containing a single PAM next to the partial target. Y axis represents total number of counts.
C) Example FRET traces from single-target multiple-PAM experiments. Top trace shows an example of a single-step binding event and a binding event that starts at a lower FRET state before transitioning to productive binding high-FRET state. Bottom trace shows an example of FRET fluctuations between the high-FRET state and lower FRET states.
D) A histogram showing the percentage of dynamic. The percentage values were obtained from five experiments on five different days. Error bars represent standard error of the mean. The percentage of events showing dynamic binding behaviour was calculated by dividing the number of binding events that showed such behaviour by the total number of binding events.
E) A scatter plot showing the time Cas9 spends in a lower FRET state before transitioning to a high-FRET state. The dwelltimes increase with the number of PAMs adjacent to the target. Such dwelltime could not be determined for a construct containing a single PAM. The values were obtained from five experiments on five different days. Error bars represent standard error of the mean.

Binding to DNA containing a single PAM next to the partial target resulted in single-step events showing a stable expected high FRET efficiency of 0.96 (Fig. 3b, c). However, increasing number of adjacent PAM sites next to the target displayed an increasing percentage of binding events that either start at a lower FRET state before transitioning to the productive binding FRET state or show fluctuations between a clearly defined high FRET state and various lower FRET states (Fig 3c, d). In particular, the percentage of events that show either fluctuations or 2-step binding (“dynamic events”) shows a 6-fold increase from 2.03 ± 1.61% for the 1xPAM construct to 13.9 ± 2.51% for the 9xPAM construct (Fig. 3d). Furthermore, the time Cas9 spends in an initial low FRET state before transitioning to high FRET state (Δτ_t_), which indicates on-target binding, was found to increase with increasing number of PAM sequences adjacent to the target (Fig. 3e). Together with the observation of the increasing dwelltime when the number of neighboring PAM sites increases in Fig. 2d, these results further suggest that Cas9 not only uses 3-D diffusion alone during target search, but can also find a target site by laterally probing neighboring PAM sites. Therefore, upon failing to form a stable R-loop Cas9 does not necessarily dissociate from the DNA strand but can go back to scanning the PAM sequences in a 1- dimensional fashion.

### Mechanism of Lateral Diffusion

The observations of increasing τ_2_ with increasing number of neighboring PAM sequences together with the observed transitions from lower FRET state to high FRET state when a partial target was present suggest that lateral diffusion may indeed aid Cas9 target search. In order to investigate this mechanism more systematically, we designed tandem target DNA constructs, where identical partial targets were placed at different distances: 6, 9, 12, and 23bp (Fig. 4a). In this assay, binding to one target (H) would yield a high FRET value and binding to the second target (L) would correspond to a lower value (Fig. 4a). We used a partial match (3 nt) between crRNA and DNA target to investigate how the presence of a second target site influences Cas9 binding dynamics and whether it can laterally diffuse between the two target sites. Control experiments with a single target at each distance showed no fluctuations in FRET efficiency (**Supplementary Fig. 3a**).

**Figure 4.**
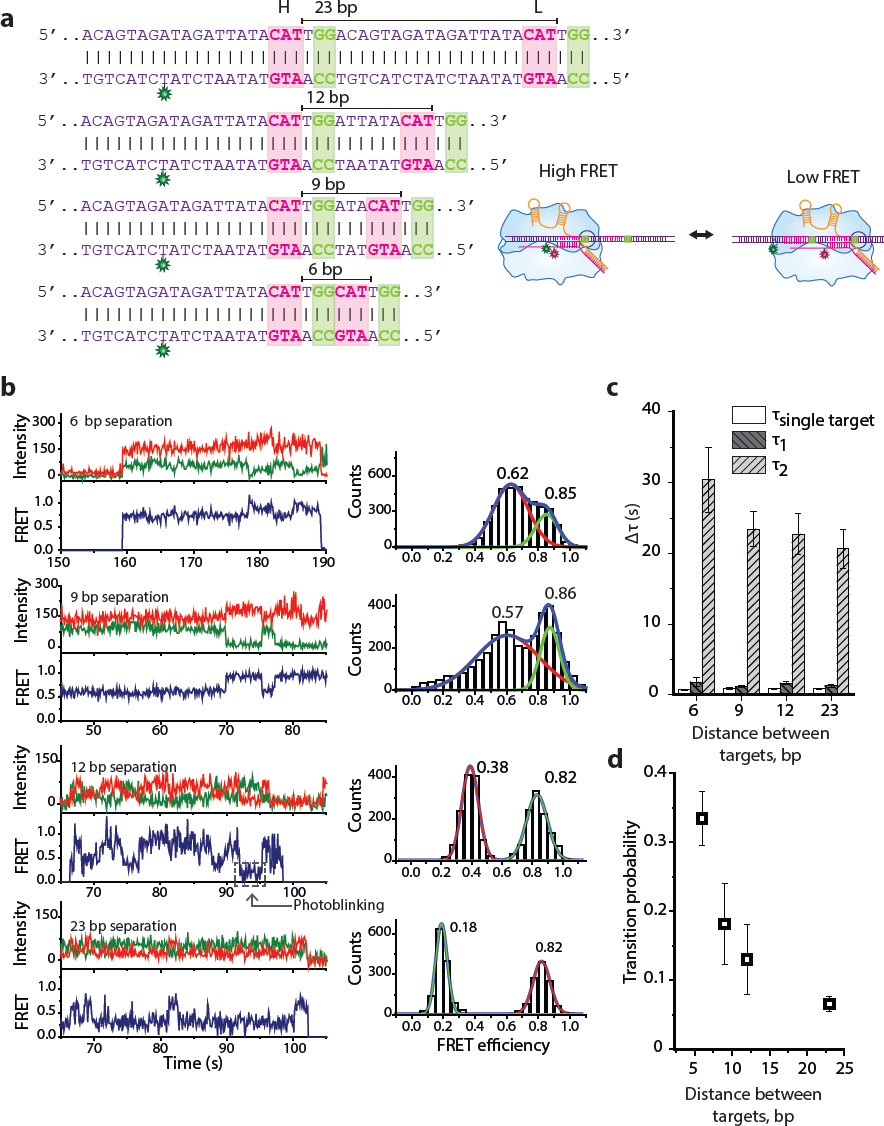
Mechanism of lateral diffusion. A) DNA sequences used in tandem target experiments. Complementary sequences to crRNA are marked in pink boxes and PAMs are marked in green boxes. Distances between the protospacers are indicated above the sequences. An illustration of Cas9 binding to each target site resulting in either high FRET (target H) or low FRET (target L) is shown on the right.
B) Example fluorescence traces showing Cas9 transitioning between H and L targets with increasing target distances (right). FRET histograms constructed from events that show fluctuations showing two FRET peaks corresponding to binding to either target. High FRET peak stays the same in each histogram, but lower FRET peak moves to lower FRET values as the distance between targets increases.
C) A histogram showing all dwelltimes from single-target controls and tandem-target experiments. τ_1_ is similar for single-target control and tandem target experiments. τ_2_ only appears in tandem target experiments and is over 20-fold higher than τ_1_ or T_st_. The values represent the mean dwelltimes that were obtained by fitting dwelltime distributions by either single or double exponential decay function from five measurements. Error bars represent standard error of the mean.
D) A scatter plot showing the probability of Cas9 transitioning between targets before dissociation. The values are averages from five measurements and error bars represent standard error of the mean. The transition probability *(p = T /* (*T+D*)) was calculated by summing up all the number of FRET transitions (T) and dividing *T* by the sum of *T* and the total number of dissociation events (D).

In the tandem target assay, Cas9 was directly observed to switch between two FRET states in a single binding event for each target separation (Fig. 4b). The observed two FRET peaks in histograms from transition events agree with the values from single target controls (Fig. 4b, **Supplementary Fig. 3a**), confirming that the fluctuations are arising due to Cas9 shuttling between two target sites. The probability of translocating to a neighboring target before dissociation that arose due to 1-D diffusion was highest at ∼0.35 when the distance between protospacers was 6 bp (Fig. 4d). At 9-bp and 12-bp separation the probability dropped to ∼0.19 and ∼0.13, respectively. At 23-bp distance, the probability dropped to ∼0.07 - a 5-fold decrease compared to 6-bp separation.

The measured dwelltimes for a single target at each distance from the dye (Fig. 4c, **supplementary figure 3b**) followed a single exponential decay with dwelltimes (τ_st_) lower than 1 s which is in agreement with literature ^27^. In contrast, the dwelltime distribution for tandem target constructs was characterized by a double exponential decay (Fig. 4c, **supplementary figure 3c**). Short binding events with a similar dwelltime as single target controls were observed with all constructs regardless of the distance between protospacers ( τ_1_). However, a second type of events observed had strikingly long dwelltimes (τ_2_) of 20-30 s (Fig. 4c). The presence of such binding events with more than a 20-fold increase in dwelltime compared to a single-target suggests that Cas9 is experiencing a strong synergistic effect when two targets are neighboring each other.

## Discussion

As a means of prokaryotic defense against invading foreign genetic elements, Cas9 has to be able to find its target in a crowded cellular environment, among kilo-bases of DNA. The target search becomes even more complicated when Cas9 is applied in eukaryotic cells as a genome engineering tool ^15,18^. In such situations where a protein needs to sample a myriad of sequences before finding a cognate target, facilitated diffusion has been shown to speed up target search as opposed to 3-dimensional diffusion alone ^26,28–33^. We propose that, once Cas9 finds a PAM sequence by 3-D collisions, it is able to diffuse laterally on a DNA strand. By competing with the dissociation process, this lateral diffusion mode might aid PAM finding and consequently target recognition. We determined that lateral diffusion of Cas9 primarily occurs in a local manner of ∼20 basepairs, when search for PAM and partial complementarity between DNA and crRNA. This explains the disagreement with previous studies which suggested lateral diffusion does not occur in Cas9 target search, since such distances could not be investigated due to the diffraction limit of other microscopy techniques ^20^.

We speculate that a limiting factor for Cas9 diffusion may not only be distance, but also the need to open the DNA duplex which is energetically unfavorable if a protein without a helicase domain were to laterally diffuse long distances. Structural data showed that Cas9 interacts with PAM sites directly, without opening up the double-stranded structure or the involvement of DNA-RNA interactions ^34^. Thereby, the lateral diffusion for PAM search would be more effective than for PAM and partial complementarity. This speculation is supported by our observation that multiple neighboring PAM sites provide a binding site for Cas9 and allow Cas9 to interrogate an adjacent target site. This observation is in contrast to rapid dissociation from a PAM when an adjacent target is not present ^27^.

Based on our findings we propose a model in which PAM sequences drive lateral diffusion as the protein directly interacts with them, as shown by structural studies (Fig. 5) ^34^. If upon binding to a PAM site a matching target is not found, Cas9 can dissociate or diffuse locally on the DNA strand until another PAM site is found. If a matching DNA sequences flanks the PAM, Cas9 checks for complementarity and if it is not sufficient for stable binding, it can dissociate or again, diffuse laterally until another PAM is found, as shown by our tandem target assays. Such a process repeats until a target with a sufficiently high degree of complementarity (9-12nt) is found and Cas9 cannot further dissociate. Therefore, we expand the knowledge of Cas9 target search mechanism by showing that it is a combination of 3-dimensional and 1-dimensional diffusion along the DNA strand and that the PAM sequences are not only important as the first step of target recognition but also drive target search.

**Figure 5.**
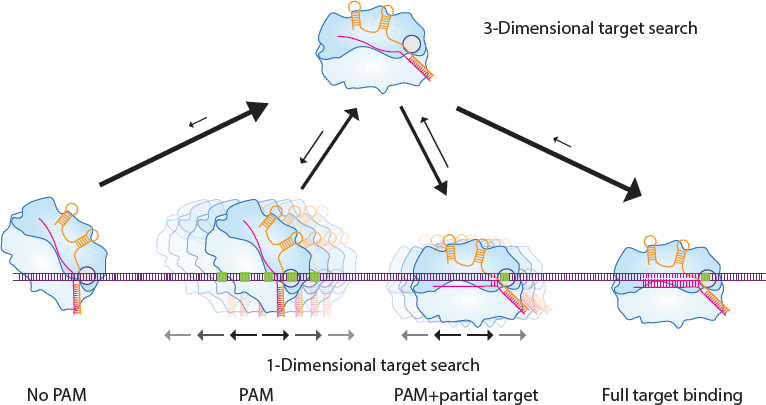
Model for Cas9 target search. Upon binding to DNA without PAM, Cas9 rapidly dissociates. Upon binding to a PAM Cas9 can dissociate or diffuse locally until another PAM site is found. If a matching DNA sequences flanks the PAM, Cas9 checks for complementarity and if it is not sufficient for stable binding, it can dissociate or diffuse laterally until another PAM is found. Such a process repeats until a target with a high enough degree of complementarity (9-12nt) is found and Cas9 cannot further dissociate.

In order to increase efficiency in genome editing, an in-depth knowledge of how Cas9 finds its target is crucial. Therefore, our results will be important when choosing target sites during genome editing. Neighboring PAM sites close to a desired target will cause the protein to stay bound to DNA for longer, thereby increasing the chance of finding the correct target faster. On the contrary, if no target is present next to a PAM-rich DNA site, such PAM clusters could be used as decoy binding sites by phages in order to prevent Cas9 binding to a cognate target during Cas9 DNA interference. In addition to importance in genetic engineering, our results suggest that the strong interaction and lateral diffusion between PAM sites could be important in bacterial defense against phages. Cas9 has been shown to be important in recognizing the PAM during the CRISPR adaptation step, together with Cas1-Cas2-Csn2 complex ^35^. Therefore, PAM density in the invader’s genome could potentially play a role in selecting which targets will be integrated in the CRISPR locus.

## ACKNOWLEDGEMENTS

We would like to acknowledge Luuk Loeff and Sungchul Kim for their assistance in setting up the experiments. We are grateful to Chun Heung Wong, Iasonas Katechis and Sabina Colombo for critical reading of the manuscript. C.J. was funded NWO (Netherlands Organisation for Scientific Research) VIDI grant (864.11.005) and the Frontiers of Nanoscience program (NWO). J.-S.K. was supported by grants from the Institute for Basic Science (IBS-R021-D1).

## AUTHOR CONTRIBUTIONS

V.G., S.H.L. performed single-molecule experiments; V. G., S. H. L performed data analysis; S. H. L, and T. B. performed protein purification; V.G., S.H.L., C.J. and J.S.K. wrote and discussed the manuscript.

## MATERIAL AND METHODS

### Recombinant SpCas9 purification

The pET plasmid encoding (6x)His-tagged Cas9 was transformed into BL21 (DE3), Rosetta. Transformed bacterial cells were moved to a 400ml of fresh LB medium containing 50ug/ml kanamycin. Incubate the culture with shaking (200rpm) at 18°C for 24 hours. Optical density was monitored and Cas9 protein expression was induced (A550=0.6) by using 0.5mM IPTG at 18C for 24hours. After the cells were harvested by by centrifugation (5000xg) for 10minutes (at 4°C), bacterial cells were resuspended with lysis buffer [20 mM Tris–HCl (pH 8.0), 400mM NaCl, 10mM b-mercaptoethanol, 1% Triton X-100, 50mg aprotinin, 50mg antipain, 50mg bestatin, 1mM PMSF (phenylmethylsulfonyl fluoride)] (Sigma-Aldrich) and sonicated on ice. The lysate was centrifuged at 6000 rcf for 10min(4°C) and supernatant solution was mixed with 2ml of Ni-NTA slurry (Qiagen) at 4°C for 1 and half hour. The lysate/Ni-NTA mixture was loaded onto a column (Biorad) with capped bottom outlet. Loaded sample was washed multiple times with pre-made wash buffer [20 mM Tris–HCl (pH 8.0), 400mM NaCl, 10mM b-mercaptoethanol] and (6x)His-tagged SpCas9 was eluted with Elution Buffer [20 mM Tris–HCl (pH 8.0), 400mM NaCl, 10mM b-mercaptoethanol, 200mM Imidazole] (**Supplementary Fig. 1a**). Finally, buffer containing eluted SpCas9 protein was changed to storage buffer[10mM HEPES-KOH (pH 7.5), 250mM KCl, 1mM MgCl2, 0.1mM EDTA, 7mM b-mercaptoethanol and 20% glycerol] by using centrifugal filter (Amicon Ultra 100K). The purified SpCas9 protein was freezed with liquid nitrogen and stored at −80°C.

### Biotinylation of the recombinant SpCas9

The process of linking biotin to the recombinant protein was carried out in-vitro and proceeded during the process of protein purification. After loading the SpCas9 over-expressed bacterial lysate and Ni-NTA mixture onto a column (Biorad), mixed sample was washed multiple times with wash buffer [20 mM Tris–HCl (pH 8.0), 400mM NaCl]. Then we added 10 fold molar excess of maleimide-biotin (Sigma-Aldrich) to SpCas9 solution and incubate for overnight at 4°C (mix gently with rotator). To get rid of unbound maleimide-biotin chemicals, mixed sample was washed sufficiently with wash buffer [20 mM Tris–HCl (pH 8.0), 400mM NaCl]. Finally, biotinylated SpCas9 protein was eluted with elution Buffer [20 mM Tris–HCl (pH 8.0), 400mM NaCl, 200mM Imidazole], then the protein concentration was measured by spectrophotometer (Nanodrop 2000, Thermo Fisher Scientific). Eluted SpCas9 protein was further purified with size exclusion chromatography. The biotinylation degree of the wild-type SpCas9 protein was calculated with commercial kit (Pierce) and it reached about 100% for two Cysteine site (Cys80/Cys574). Biotinylated SpCas9 protein was storaged in storage buffer [10mM HEPES-KOH (pH 7.5), 250mM KCl, 1mM MgCl2, 0.1mM EDTA, 7mM b-mercaptoethanol and 20% glycerol] and purified protein was freezed in liquid nitrogen and store at −80°C.

### Preparation of the single-guide RNA

We used In-vitro RNA transcription (DNA template, T7 RNA polymerase (NEB) 5ul, 10x Buffer 10ul, rNTP mix (2.5mM each), MgCl2 10mM, DTT 1mM, H2O up to 100ul, RNAse inhibitor (NEB) 0.5ul, Total 100ul reaction) to generate single-guide RNAs. DNA template contains X20 target protospacer sequence which is complementary to the RNA strand. After RNA transcription, DNA template was removed by DNase (NEB) treatment. Then pure single-guide RNA was purified and concentration was measured by spectrophotometer (Nanodrop 2000, Thermo Fisher Scientific).

### In-vitro DNA cleavage assay with wild-type and biotinylated SpCas9

In-vitro cleavage experiments were performed with sgRNA and SpCas9 proteins purified at high purity. DNA containing the target site was prepared by PCR, and higher molar concentration of the SpCas9 protein and biotinylated SpCas9 was treated at the same molarity. The sgRNA was added at a molar ratio three times greater than the protein (final molar ratio, DNA: protein: sgRNA = 1:3:9) with a complementary sequence to the target site. Target DNA, SpCas9 protein and sgRNA were mixed and incubated at 37°C for 1hour. The cleaved DNA product was separated on the 1.5% agarose gel and cleavage ratio was calculated by ImageJ software.

### Single-molecule two-color FRET

Single-molecule fluorescence measurements were performed with a prism-type total internal reflection fluorescence microscope. 0.1mg/ml Streptavidin was added to a polyethylene glycol-coated quarts surface and incubated for 2 minutes before being washed with T50 (10 mM Tris–HCl (pH 8.0), 50 mM NaCl). Biotinylated Cas9 was pre-incubated with Cy5-labeled crRNA and tracrRNA (ratio 1:2:4) at 37 degrees for 20 minutes in NEB buffer 3 (100 mM NaCl, 50 mM Tris–HCl, 10 mM MgCl2, 1 mM DTT) and then added to the chamber containing Streptavidin. After 2 minute incubation unbound Cas9 and RNA molecules were washed away with an imaging buffer (50mM HEPES-NaOH [pH7.5], 10mM NaCl, 2mM MgCl2, 1% glucose (Dextrose monohydrate), 1mM Trolox (2.5mg/10ml), 1mg/ml glucose oxydase [Sigma], 170ug/ml catalase [Merck]). 8 nM Cy3 labeled DNA substrate in imaging buffer was added to the channel. A reference video of immobilized Cy5-labeled Cas9:RNA complexes were made. Following the reference video, Cy3-labeled DNA molecules were excited using a 532 nm diode laser. Fluorescence signals of Cy3 and Cy5 were collected through a 60× water immersion objective (UplanSApo, Olympus) with an inverted microscope (IX73, Olympus). The 532 nm laser scattering was blocked out by a 532 nm long pass filter (LPD01-532RU-25, Semrock). The Cy3 and Cy5 signals were separated with a dichroic mirror (635 dcxr, Chroma) and imaged using an EM-CCD camera (iXon Ultra, DU-897U-CS0-#BV, Andor Technology).

### Data acquisition and analysis

Using a custom-made program written in Visual C++ (Microsoft), a series of CCD images of time resolution 0.1s was recorded. The time traces were extracted from the CCD image series using IDL (ITT Visual Information Solution) employing an algorithm that looked for fluorescence spots with a defined Gaussian profile and with signals above the average of the background signals. Colocalization between Cy3 and Cy5 signals was carried out with a custom-made mapping algorithm written in IDL. The extracted time traces were processed using Matlab (MathWorks) and Origin (Origin Lab).

## References

1. Barrangou, R. et al. CRISPR provides acquired resistance against viruses in prokaryotes. Science 315, 1709–12 (2007).

2. Marraffini, L.A. CRISPR-Cas immunity in prokaryotes. Nature 526, 55–61 (2015).

3. Makarova, K.S., Grishin, N.V., Shabalina, S.A., Wolf, Y.I. & Koonin, E.V. A putative RNA-interference-based immune system in prokaryotes: computational analysis of the predicted enzymatic machinery, functional analogies with eukaryotic RNAi, and hypothetical mechanisms of action. Biol Direct 1, 7 (2006).

4. Makarova, K.S. et al. An updated evolutionary classification of CRISPR-Cas systems. Nat Rev Microbiol 13, 722–36 (2015).

5. Mohanraju, P. et al. Diverse evolutionary roots and mechanistic variations of the CRISPR-Cas systems. Science 353, aad5147 (2016).

6. Wiedenheft, B., Sternberg, S.H. & Doudna, J.A. RNA-guided genetic silencing systems in bacteria and archaea. Nature 482, 331–8 (2012).

7. Amitai, G. & Sorek, R. CRISPR-Cas adaptation: insights into the mechanism of action. Nat Rev Microbiol 14, 67–76 (2016).

8. Bolotin, A., Quinquis, B., Sorokin, A. & Ehrlich, S.D. Clustered regularly interspaced short palindrome repeats (CRISPRs) have spacers of extrachromosomal origin. Microbiology 151, 2551–61 (2005).

9. Mojica, F.J., Diez-Villasenor, C., Garcia-Martinez, J. & Soria, E. Intervening sequences of regularly spaced prokaryotic repeats derive from foreign genetic elements. J Mol Evol 60, 174–82 (2005).

10. van der Oost, J., Westra, E.R., Jackson, R.N. & Wiedenheft, B. Unravelling the structural and mechanistic basis of CRISPR-Cas systems. Nat Rev Microbiol 12, 479–92 (2014).

11. Brouns, S.J. et al. Small CRISPR RNAs guide antiviral defense in prokaryotes. Science 321, 960–4 (2008).

12. Jinek, M. et al. A programmable dual-RNA-guided DNA endonuclease in adaptive bacterial immunity. Science 337, 816–21 (2012).

13. Gasiunas, G., Barrangou, R., Horvath, P. & Siksnys, V. Cas9-crRNA ribonucleoprotein complex mediates specific DNA cleavage for adaptive immunity in bacteria. Proc Natl Acad Sci U S A 109, E2579–86 (2012).

14. Deltcheva, E. et al. CRISPR RNA maturation by trans-encoded small RNA and host factor RNase III. Nature 471, 602–7 (2011).

15. Hsu, P.D., Lander, E.S. & Zhang, F. Development and applications of CRISPR-Cas9 for genome engineering. Cell 157, 1262–78 (2014).

16. Jiang, W., Bikard, D., Cox, D., Zhang, F. & Marraffini, L.A. RNA-guided editing of bacterial genomes using CRISPR-Cas systems. Nat Biotechnol 31, 233–9 (2013).

17. Barrangou, R. & Doudna, J.A. Applications of CRISPR technologies in research and beyond. Nat Biotechnol 34, 933–941 (2016).

18. Cong, L. et al. Multiplex genome engineering using CRISPR/Cas systems. Science 339, 819–23 (2013).

19. Jones, D.L. et al. Kinetics of dCas9 target search in Escherichia coli. Science 357, 1420–1424 (2017).

20. Sternberg, S.H., Redding, S., Jinek, M., Greene, E.C. & Doudna, J.A. DNA interrogation by the CRISPR RNA-guided endonuclease Cas9. Nature 507, 62–7 (2014).

21. Chandradoss, S.D. et al. Surface passivation for single-molecule protein studies. J Vis Exp (2014).

22. Brown, M.W. et al. Assembly and translocation of a CRISPR-Cas primed acquisition complex. bioRxiv (2017).

23. Szczelkun, M.D. et al. Direct observation of R-loop formation by single RNA-guided Cas9 and Cascade effector complexes. Proc Natl Acad Sci U S A 111, 9798–803 (2014).

24. Chandradoss, S.D., Schirle, N.T., Szczepaniak, M., MacRae, I.J. & Joo, C. A Dynamic Search Process Underlies MicroRNA Targeting. Cell 162, 96–107 (2015).

25. Grimson, A. et al. MicroRNA targeting specificity in mammals: determinants beyond seed pairing. Mol Cell 27, 91–105 (2007).

26. Berg, O.G., Winter, R.B. & von Hippel, P.H. Diffusion-driven mechanisms of protein translocation on nucleic acids. 1. Models and theory. Biochemistry 20, 6929–48 (1981).

27. Singh, D., Sternberg, S.H., Fei, J., Doudna, J.A. & Ha, T. Real-time observation of DNA recognition and rejection by the RNA-guided endonuclease Cas9. Nat Commun 7, 12778 (2016).

28. Halford, S.E. & Marko, J.F. How do site-specific DNA-binding proteins find their targets? Nucleic Acids Res 32, 3040–52 (2004).

29. Riggs, A.D., Bourgeois, S. & Cohn, M. The lac repressor-operator interaction. 3. Kinetic studies. J Mol Biol 53, 401–17 (1970).

30. Hammar, P. et al. The lac repressor displays facilitated diffusion in living cells. Science 336, 1595–8 (2012).

31. Leith, J.S. et al. Sequence-dependent sliding kinetics of p53. Proc Natl Acad Sci U S A 109, 16552–7 (2012).

32. Gorman, J. et al. Single-molecule imaging reveals target-search mechanisms during DNA mismatch repair. Proc Natl Acad Sci U S A 109, E3074–83 (2012).

33. Ragunathan, K., Liu, C. & Ha, T. RecA filament sliding on DNA facilitates homology search. Elife 1, e00067 (2012).

34. Anders, C., Niewoehner, O., Duerst, A. & Jinek, M. Structural basis of PAM-dependent target DNA recognition by the Cas9 endonuclease. Nature 513, 569–73 (2014).

35. Heler, R. et al. Cas9 specifies functional viral targets during CRISPR-Cas adaptation. Nature 519, 199–202 (2015).

